# Miniaturized Device for Assessing Calcification Propensity of Biohybrid Implants Under Continuous Flow

**DOI:** 10.64898/2025.12.15.693933

**Authors:** Aaron D. Morgan, Robert Dzhanaev, Andrea Gorgels, Jan Ritter, Johanna C. Clauser, Felix Stockmeier, Lucas Stüwe, Christian Böhm, Stefan Jockenhoevel, Ulrich Steinseifer, Willi Jahnen-Dechent

## Abstract

Biohybrid implants are a promising development for cardiovascular disease treatment but suffer from problems like thrombogenesis and calcification. However, testing and validating biohybrid implants can be difficult and expensive due to material handling, fabrication methods and specialty medium components. The devices used to test potential samples can be large and expensive, requiring significant amounts of cell culture medium to operate. Additional, conventional static cell culture conditions do not accurately represent the vascular environment as shear and mechanical forces play key roles in the development of calcification. To address these challenges, a miniaturized, dual-channel flow chamber was designed and validated that allowed for real-time visualization of biohybrid calcification in a physiological environment. Computational Fluid Dynamics simulations were performed to determine the flow characteristics that generated physiological shear stress homogeneously across the sample surface. Micro particle tracking velocimetry measurements validated the simulated shear stresses near the sample surface. Two implant materials used for biohybrid construction, bovine pericardium and polycarbonate urethane, were inserted in the device and exposed to a flowing calcification medium for 14 days. Fluorescent fetuin-A was introduced into the calcification medium for real-time calcification monitoring. The two materials were compared with matched samples calcified in a large fatigue tester for 14 days. Our results showed similar material calcification for bovine pericardium and no calcification for polycarbonate urethane in the large fatigue tester and in our newly developed device. Biohybrid textile-reinforced fibrin-based scaffold populated with vascular smooth muscle cells started to calcify over 7 days in calcification medium. We conclude that this platform will provide novel insights into the origin and progression of pathological calcification and its potentially harmful health effects, which can occur as a result of tissue or metabolic abnormalities, disease, or implantation of certain biomaterials, by providing the ability to monitor the progression of calcification in biohybrid implants in real time, while also minimizing the cost and size of samples and reagents required for testing.

## Introduction

Cardiovascular calcification is a common degenerative pathology treated by replacing affected vessel parts or heart valves with artificial cardiovascular implants. The major disadvantages of cardiovascular implants currently being used are the need for lifelong anticoagulation^[1–3]^ and early implant failure due to implant calcification.^[4–7]^ Biohybrid implants combine cellular and scaffold components to overcome limitations of purely technical replacements. Bioengineered vessels, for example, provide important insights into the effect that mechanical forces have on calcification.^[8]^ However, testing and validating biohybrid implants remain a challenge for researchers as these materials, in addition to medical materials, have live cells. Their production lot size is nominally “1”, theoretically requiring damage-free and online testing of every single product. This makes manufacturing and testing complex, one reason why it is challenging to manufacture biohybrid constructs at a large scale.

Current testing devices require liter amounts of medium as well as physically large material samples.^[9, 10]^ The long times and the large volumes of fluids required for testing preclude the testing of small experimental material samples that may be in short supply, as well as the use of complex cell-compatible test environments including growth factors and nutrients that must be kept intact and sterile. Calcification testing of acellular bioprosthetic heart valves is commonly performed using a large two-chamber testing device. The material samples are exposed to a simulated body fluid (SBF), which is not cell compatible. We routinely tested valves in a large Cardiovascular Engineering Fatigue Tester, version 2, (CVE-FT2) using simulated body fluid at 300+ bpm over 2-9 weeks.^[11]^ In contrast, testing of biohybrid implants requires cell compatible calcification fluids based on cell culture medium and physiologically appropriate temperature and fluid characteristics including shear stress.

It is known that physiological shear stress is necessary to maintain healthy and effective arterial architecture.^[12–16]^ Exposure to low or unphysiological shear stress causes a loss of barrier integrity, exposing matrix and cells below.^[17]^ Loss of barrier integrity is also shown to promote extravasation of circulating cells into the subluminal matrix.^[18]^ To ensure a robust barrier, the flow must be laminar and provide homogenous shear stress to the material surface. An arterial surface must be exposed to 1-20 dyne/cm^2^ to maintain a robust, intact architecture.^[19–22]^ Given the importance of flow in the maintenance of vascular architecture, continuous flow of a cell compatible medium, as opposed to repetitive medium exchanges, is preferred.

An ideal testing system for biohybrid implants should also heed recent biomineralization research demonstrating the importance of fluid phase saturation and stability in biomimetic extracellular matrix mineralization.^[23]^ Protein-mineral complexes have been described that increase the mineral saturation and availability in calcification fluids, while preserving stability and cell compatibility. Calcium phosphate is the predominant mineral phase in vertebrate tissue calcification, and mineral-rich fluids are naturally stabilized as colloidal complexes with blood proteins fetuin-A and albumin.^[24]^ By formulating serum-containing cell culture media that contain highly saturated, yet stable mineral phases, robust cell- and material-associated calcification is produced, while artefactual spontaneous mineral precipitation from solution onto cell and material surfaces is prevented. One particularly powerful imaging reagent for monitoring calcification in live cell culture is fluorescent fetuin-A.^[25]^ Fetuin-A is a liver derived plasma protein, which serves as a mineral chaperone by binding and stabilizing calcium phosphate mineral clusters ^[26, 27]^, making it suitable for the detection of early forms of calcification.

To further advance the testing of biohybrid materials in simulated physiological environments, we have developed a novel, miniaturized, two-chamber flow device. This study aims to provide the same calcification testing capabilities to the established CVE-FT2 ^[11]^ while allowing for the use of much smaller cellularized material samples and substantially reduced volumes of fluid, facilitating the exploration of advanced biohybrid implant materials and more complex and costly calcification fluids, as well as imaging reagents. This flow chamber creates possibilities for novel calcification studies across a wide range of natural, biohybrid, and artificial implant materials while simultaneously providing the possibility for high temporal resolution through live imaging, as well as more advanced high-resolution microscopy with techniques such as long-distance two-photon microscopy.

## Methods and Materials

### Cell culture in static and flow conditions

Immortalized vascular smooth muscle cells (IM1) (kindly provided by Claudia Goettsch, RWTH Aachen University Hospital)^[28]^, iPSC-derived vascular smooth muscle cells (iVSMC, kindly provided by Cengiz Akbulut and Leon Shurgers, Univ. of Maastricht), and human kidney epithelial cells (HK-2, kindly provided by Prof. Peter Boor, RWTH Aachen University Hospital) were seeded into 24-well plates at a density of 10,000 cells per well and cultured in calcification medium CM for 7 and 21 days, respectively. CM comprised DMEM (31053-044, Lot: 203043, Gibco) with 10% FBS (P30-3033, Lot: P231101, Pan Biotech), 1% PenStrep (15140-122, Gibco), 4.1 mM total calcium and 3.0 mM total phosphate. Calcification was monitored using a live imaging station (Cellcyte X, Cytena). Fluorescent fetuin-A was added to the culture medium at 100 ug/mL to monitor calcification. C.Live Tox Green dead stain (CY.CL.KIT.001, Cytena) was used to quantify dead cells. To study the effect of flow vs. static culture with medium exchange on calcification, iVSMC were seeded into a flow chip (Ibidi 80186) and exposed to 10 dyne/cm^2^ shear stress by continuous flow for 21 days. A syringe pump (Ibidi10902) was used to produce the continuous flow. The selection of cell lines was based on physiological relevance and experimental practicality. Immortalized vascular smooth muscle cells (IM1 and iVSMC) were chosen for their established role in vascular calcification. Their rapid proliferation in static culture limited experiment duration to 7 days to prevent overgrowth. To enable longer-term static culture studies (14 days) for observing later calcification time points, HK-2 human kidney epithelial cells were chosen due to their slower proliferation rate, while still being a relevant model for pathological calcification.

### Fluorescence quantification

To simultaneously quantify the amount of calcification and cell death, fluorescent fetuin-A-mRuby3 and C.Live Tox Green were added to culture medium as above. Using the Cellcyte X live imaging system, cell cultures were monitored and fluorescence images were acquired in the red and green channels. The acquired frames were analyzed using the Cellcyte Studio software (Cytena). For calcification, background fluorescence with fetuin-A-mRuby3 in the medium was recorded at the onset of the experiment. An increase in fluorescence beyond the background threshold was interpreted by the software as calcification. The total calcified area was reported in percentage for each acquired frame at each time point. For cell death, background fluorescence was established at the onset of the experiment. Afterwards, acquired frames were segmented by fluorescence intensity against background and the number of circular fluorescent objects was counted. The total number of counted objects was reported as number of dead cells for each acquired frame at each time point.

### Chip material, fabrication, and post-fabrication processing to prepare for use

The flow chip developed in this work was comprised of acrylic, and the features of each chip were milled. The geometry of each channel was designed such that the appropriate flow characteristics were produced when both channels had equal rates of volumetric flow, allowing one or more of the devices to be driven by a single pump. This design also allowed for the pairing of different chip pieces to accommodate various thicknesses of material samples and desired flow conditions. The device maintained its internal chamber-to-chamber seal based on the thickness of the material sample and inner sealing rings, while both chambers were sealed from the outside environment by an outer seal. This allowed for sterility to be maintained in the case of sample rupture or degradation. Seals were placed in the grooves in each chip, then the chip pieces were assembled using threaded bolts. These bolts applied light sealing pressure across the chip to ensure a water-tight seal and firmly secured the sample in place.

### Establishing and maintaining controlled volumetric flow

A peristaltic roller pump was used to provide 12 ml/min volumetric flow rate of medium drawn from a closed-loop reservoir. Silicone tubing and polymer tubing connectors were used to connect the chip to the pump and reservoirs. Each of the two flow channels in the assembled chip were perfused by two reservoirs with filtered gas perfusion possible at the liquid interface of each reservoir. Each reservoir was filled with 30 mL of cell medium, holding up to 50 mL. The pump had multiple channels, allowing for the use of multiple flow chips in parallel. The pump also allowed for the removal of individual chips from the system without needing to stop the flow for other chips on the same pump.

### Computational fluid dynamic simulation for the high-flow channel

CFD simulations were performed using Autodesk CFD 2019. The channel models were imported, with “acrylic” assigned as the body material and “water” assigned to the fluid volume. The simulation environment was set to atmospheric pressure (101,325 Pascal) and body temperature (37°C). For each model, inlet and outlet boundary planes were defined to set a specific volumetric flow rate (in cm³/min), and the flow was defined as steady state. The flow was not assumed to be fully developed before entering the channel. Initial fluid velocity conditions were set to 10 cm/s.

Each model was first meshed using the default recommended settings. Then, a localized mesh modification was applied specifically to sample surface using a fine mesh density of 0.025. This significantly increased the mesh density near the sample surface, where the flow conditions were the primary concern, without unnecessarily increasing density in areas far from the sample surface. This localized approach allowed for much faster simulation iterations compared to a uniformly dense mesh. Heat transfer simulation was disabled, as no significant temperature gradient or heat generation was expected. The maximum number of simulation iterations was set to 300, and for consistency, Automatic Mesh Adaptation was not used. All other solver settings were set to default.

### Post-processing: shear stress analysis across sample surface

After the simulations were completed, the shear stress across the sample for each scenario was analyzed. To visualize and quantify the shear stresses, a result plane was aligned to the mounted sample surface (Figure 2B). Sampling lines were then placed across this surface to generate shear stress profiles. Two hundred equally spaced data points were taken from each sampling line. Five sampling lines were placed across the width of the sample (relative to the flow direction), and three were placed along its length (parallel to the flow direction). Both the result plane and the sampling lines were assigned as “summary values” in the software, enabling comparison across all simulated scenarios. Using the software’s Decision Center, the shear profiles from each sampling line were superimposed onto a single figure and exported as a data table for analysis in MATLAB.

### Post-processing: velocity analysis for flow development

To assess how the flow developed between the flow guide exit and the sample surface, a separate velocity analysis was performed on the channel’s midplane. For this, eight sampling lines were placed on the midplane velocity result plane (Figure 2A) This midplane analysis was performed only for the specific scenario selected by the following comparative methodology.

### CFD results processing and scenario ranking in MATLAB

To objectively select the optimal flow chip geometry and operating condition, we developed a comparative methodology in MATLAB. Each simulated scenario (a unique combination of flow guide position and volumetric flow rate) was assigned a "deviation score." This score quantified the total deviation of the simulated shear stress across the sample surface from our target profile: a uniform shear stress of 10 dyne/cm². The scenario with the lowest deviation score, corresponding to the most homogeneous shear distribution, was selected for all experimental work. This analysis identified a flow guide position of 4 mm and a volumetric flow rate of 12 ml/min as the optimal parameters.

### MicroPTV measurements validating flow characteristics from CFD simulations

The chosen channel parameters were recreated in an acrylic flow chip. This chip was modified to be compatible with the microPTV system but was fluid dynamically identical to the design used for cell culture. The setup allowed the microPTV system to observe fluorescent tracer particles (diameter 3.2 µm) flowing through the channel. A high-frequency Nd:YAG laser illuminated the particles at a wavelength of 532nm. The resulting fluorescent signal was recorded by two high-speed cameras through a stereomicroscope. This microPTV setup enabled a recorded volume of 5.4 mm × 3.4 mm × 1.0 mm. Tracer particles were suspended in solution and pumped through the device at fixed volumetric flow rates of 2 ml/min and 10 ml/min.

The recorded images were analyzed using DaVis software. The particle images were post-processed by subtracting the time-averaged intensity for each pixel. Single particle tracks were then reconstructed using the Shake-the-Box algorithm. These tracks were subsequently transformed into a 3D velocity field with components in the x-, y-, and z-directions. Finally, these measured velocities were compared to the velocities from the corresponding CFD simulation to validate the accuracy of the simulated model.

### Bovine pericardium and polycarbonate urethane patches in calcification medium

Calcification testing of test materials was done using the newly developed flow chip and results were compared to an established system, the CVE-FT2.^[11]^ Bovine pericardium patches (SJM Pericardial Patch with EnCap Technology, St. Jude Medical) and polycarbonate urethane patches were analyzed. Patches were produced by evaporating a 5% w/w PCU (Carbothane TPU PU 3775, Lubrizol) chloroform solution. Both materials were cut into circular patches of 38 mm diameter for the CVE-FT2. Patches were clamped between two patch holders of the CVE-FT2 with an outer diameter of 38 mm and an inner diameter of 16 mm and placed into separate compartments to maintain partly closed conditions. The patches were loaded sinusoidally with physiological pressures comparable to an aortic valve with a peak pressure difference of at least 100 mm Hg for 5% of the cycle duration. The test ran at 300 bpm to facilitate an accelerated procedure. Every compartment was filled with calcification fluid from the same batch to keep conditions such as ionic concentrations equal. The temperature was set to 37 °C. To monitor stability of the calcification fluid, a sample in a PE bottle was put into the CVE-FT2 as a control.

For flow chip testing, 25 mm diameter material samples were clamped into the flow chip and exposed to calcification fluid for two weeks, with fluid exchanges every week. The incubator conditions were maintained at 37°C and 5% CO_2_. The flow rate was set to 2 ml/min, then increased by 2 ml/min every 30 minutes until a volumetric flow rate of 12 ml/min was reached. This stepwise increase in flow velocity allows time for cells to adapt to stresses introduced by the continuous flow. If the flow is initiated at 12 ml/min from rest, cells often detached or responded negatively. The fluid was changed weekly. Temperature and pH were recorded throughout the experiment.

Calcification visualization was enabled using the plasma protein fetuin-A, which binds calcium phosphate mineral.^[25]^ The bovine fetuin-A (Sigma, F-2379) was chemically labelled using Alexa Fluor 546 (A20102, Thermo Fisher Scientific GmbH, Dreieich, Germany) according to the manufacturer’s protocol. Patches were stained using fluorescence labeled fetuin-A at a final concentration of 0.1 mg/ml for 20 minutes at 37°C. Stained patches were photographed with a Leica DMI 6000 microscope.

### Calcium content of calcified material

The calcium content of calcified samples was determined using a Calcium Assay kit (Randox CA590). Samples calcified in CVE-FT2 were cut into two samples. One sample was taken whole, with calcified scaling intact. One sample had the scaly surface calcification manually removed and was taken as two samples, one of purely surface calcification scales and one of only internally calcified pericardium. The pericardium calcified in the flow chip was taken as the fourth sample. The total surface area of each sample was measured. Samples were placed into 1 ml of 0.6 mM hydrochloric acid solution and placed into an Eppendorf tube shaker overnight, heated to 37°C. The following day, 1 ml of neutralizing solution was added to the samples (2.4 g/L ammonium chloride, 5% ammonium hydroxide, in demineralized water). Dilution series were prepared for each sample and the Randox calcium assay was prepared in a 48-well plate, following the manufacturer’s protocol. Absorbance was recorded at 570 nm using a plate reader (FLUOstar Optima, BMG Labtech).

### Textile-reinforced biohybrid construct

A textile-reinforced biohybrid scaffold was created using two materials: PBN II nonwoven 136 g/m^2^ (Cerex advanced fibers, Cantonment, FL, United States) and a tubular warp-knitted structure with 22 mm diameter (Tuell-filet, 24 filaments, E15) produced from PET multifilament (78 dtex f 24, ICF Mercantile LLC, NJ, USA). The nonwoven PBN II was cut into 20×20 mm frames with inside dimensions of 16×16 mm. The mesh was set using heat on a 24×3 mm steel bar for 10 min at 200⁰C. After setting, the mesh was laser-cut into 10 mm wide loops. The nonwoven and mesh are welded together using a soldering iron. Mesh material was then cut to leave 3 rows of stitches suspended across the frame. The textile mesh was then embedded into a fibrin-based hydrogel containing VP-co-GMA copolymer.^[29–31]^ Immortalized IM3 vascular smooth muscle cells were seeded onto the scaffold at a concentration of 500,000 cells/mL in 5 mL of medium. After 48 hours, the construct was transferred to another well to remove unattached cells. Medium was exchanged every 48 hours until brightfield microscopy confirmed cell penetration and proliferation throughout the construct. Once cell density was deemed sufficient, the construct was placed into the flow chamber and exposed to ∼10 dyne/cm^2^ under continuous, unidirectional flow in calcification medium. The construct was kept under continuous flow for 7 days and then assessed inside of the assembled system before being removed from the chip for higher resolution microscopy. CellTracker Green CMFDA (Invitrogen, #C7025) was resuspended in DMSO and added to the circulating calcification medium at a concentration of 10 μM.

## Results

### Live imaging of static vs flow conditions in vascular smooth muscle cell calcification

Cells were cultured in microwell plates under static conditions and in an IBIDI flow chip under continuous flow for 7 to 21 days. To model calcification under static conditions over different timeframes, we utilized cell lines with distinct proliferation rates. Vascular smooth muscle cells (VSMCs) were used for their high physiological relevance to cardiovascular calcification, but their rapid growth constrained static experiments to 7 days. To extend observations to 14 days without cell overgrowth, we employed slower-proliferating human kidney epithelial cells (HK-2), which also provide a model for pathological calcification. This allowed longer runtimes of 14 days and provided the opportunity to observe calcification related to medium exchange over this longer time. Calcification was measured using live imaging of fluorescent fetuin-A. Cell death was measured using C.Live Tox Green, a nuclear stain marking dead cells, which is non-toxic and therefore compatible with live cell imaging. Figure 1 illustrates calcification and cell viability in static and continuous flow cell culture models of human vascular smooth muscle cells (IM-1 and iVSMC) and kidney epithelial cells (HK-2) that were chosen for their proliferation rates to match the desired culture time without overgrowth. In static cultures of IM1 cells (Figure 1A) medium exchange on day 3 triggered cell death and subsequent calcification. Cell death and calcification also increased after each medium exchange (vertical dotted lines in Figure 1B) when slow growing HK-2 cells were analyzed for two weeks instead of fast growing IM1 cells for one week. Each medium exchange also rinsed loosely attached dead cells, reducing the overall number of dead cells, while ECM-calcified cells remained attached unless forcefully rinsed with high flow (not shown). In conditions of continuous flow however, cell death and calcification of iVSMC (Figure 1C) progressed steadily over time as would be expected for a continuous process. A noticeable delay in the onset of calcification relative to the increase in cell death was observed suggesting that continuous medium flow prevented sudden deposition of mineral and detachment of dead cells observed after every medium exchange in static culture. Fluorescent fetuin-A staining (Figure 1D) showed more calcification in the center of the flow path, where shear stress was the highest. FDA staining (Figure 1E) showed that despite widespread calcification, cell viability was high, and the channel was filled with a dense, confluent cell sheet. Alizarin Red S staining (Figure 1F) showed extensive calcification throughout the channel, with the most calcification seen in areas of highest shear stress, corroborating the fetuin-A staining.

**Figure 1.**
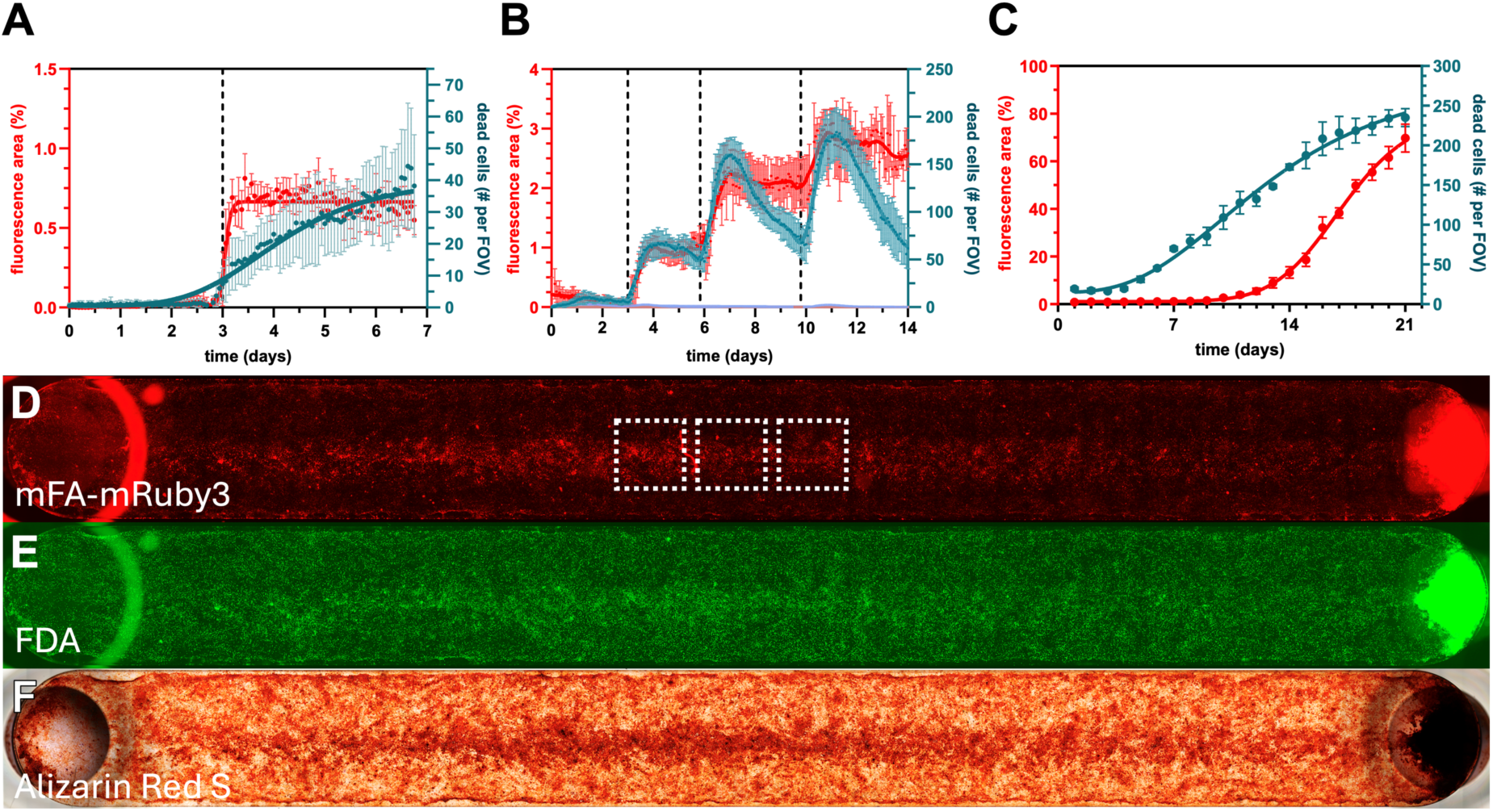
Calcification dynamics and cell viability in static and continuous flow cell culture models. Calcification and cell death were recorded using a live imaging system (Cellcyte X, Cytena). (A-C) Left axis: fetuin-A fluorescence representing calcification, given as percent of total area. Red dots with red vertical lines represent the mean ± SD of 3-6 data points as detailed below, and red solid line represents non-linear curve fit. Right axis: Dead cells stained by C.Live Tox Green, given as number of cells per field of view. Green dots with vertical lines represent the mean ±SD, and green solid lines represent non-linear data fit (A) Immortalized vascular smooth muscle cells (IM1) were kept in static culture for 7 days with one medium exchange on day 3 (black vertical dotted line). (B) Human kidney epithelial cells (HK-2) were kept in static conditions for 14 days with medium exchange on days 3, 6, and 10 (vertical dotted lines). (C): iPSC-derived vascular smooth muscle cells (iVSMC) were kept under continuous flow for 21 days. Calcification and cell death were recorded as in A and B based on live images taken from 3 fields of view (FOV) outlined with white dotted squares in D. (D-F): Microscopic tile scans of the flow chip used in C after 21 days. Distance from inlet (F, left circle) to outlet (right circle) is 50 mm. Channel width is 5 mm. (D) Red fluorescent fetuin-A. (E) Green fluorescent Fluorescein Diacetate (FDA). (F) Red Alizarin Red S. Each measurement shown in (A) comprises six measured fields of view (n = 6). Each measurement shown in (B) comprises five measured fields of view (n = 5). Each measurement shown in (C) comprises three measured fields of view (n = 3). Error bars depict standard deviation with very small errors not shown.

### Flow chip design compatible with a variety of materials and sample sizes

Figure 2 illustrates the design and performance characteristics of the proprietary flow chip, which was designed as a three-piece acrylic system (Figure 2A) to accommodate material samples of varying thickness and to enable real-time microscopy. The assembled device (Figure 2B) featured two independent fluid channels that perfuse opposite sides of the mounted sample. A key design principle was to operate both channels at the same volumetric flow rate yet generate distinct shear environments. The “high shear” channel was designed to accelerate flow and apply physiological shear stress (8-12 dyne/cm^2^) to the sample surface. In contrast, the “low shear” channel was designed to provide minimal fluid mechanical stimulation under identical flow conditions. This configuration allowed for the simultaneous investigation of material response to two different flow environments. The low-shear piece was sealed with a round glass coverslip, while the transparent top piece and a cut-out in the center piece created an unobstructed optical path through the sample.

**Figure 2.**
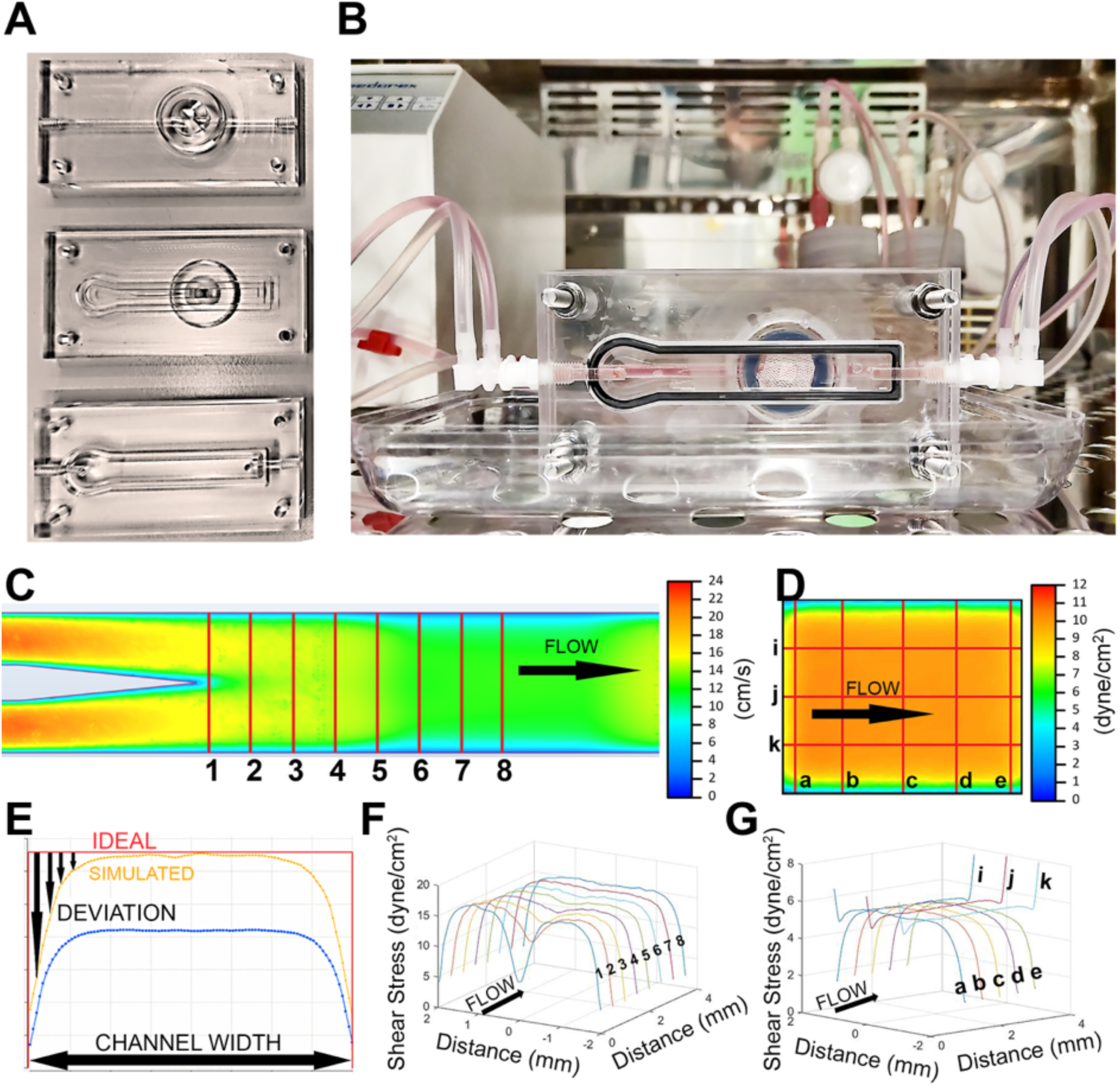
Three-piece flow chip, experimental setup, and simulation of fluid dynamics. (A) Three prepared acrylic chip pieces that, when assembled, form the high-shear channel (bottom and middle pieces) and the low-shear channel (middle and top pieces). Seals are placed in the grooves of each piece, and the sample is mounted in the chip directly over the center cut-out for microscopy. (B) Flow chip experimental setup consisting of the flow chip with a mounted sample, a peristaltic pump providing continuous flow, and two independent liquid reservoirs with filtered gas exchange. (C) Simulation results for midplane flow velocity at 12 mL/min flow rate. Direction of flow is shown. Eight sampling lines (1-8) on the midplane, perpendicular to the flow direction, are shown where velocities were recorded. Velocity is indicated by the color bar. (D) Simulated shear stress on mounted sample surface. Direction of flow is shown. Eight sampling lines (a-k), distributed across the sample surface, are shown where shear stress was recorded. Shear stress is indicated by the color bar. (E) Example of simulation scoring procedure for a single sampling line. Simulated values for shear stress (“SIMULATED”) were compared against a target value (“IDEAL” shown in red, 10 dyne/cm^2^) and the deviation from the target value (“DEVIATION” indicated by black arrows) was recorded for each point along each sampling line. The sum deviation from “ideal” was used to select the conditions that provided the most homogenous profile of shear stress on the sample surface. (F) Example of simulation of development of shear stress downstream of the flowguide. Direction of flow is shown. Sampling line 1 lies at the end of the flowguide while sampling line 8 lies on the midplane, directly above the center of the mounted sample. Midplane shear stress can be seen to develop into a homogenously distributed profile from sampling lines 6-8, indicating laminar flow. (G) Example of simulated shear stress on the mounted sample surface. Direction of flow is shown. Shear stress can be seen to develop into a homogenous shear profile on the sample surface.

### CFD analysis for determining in-stream flow guide position and volumetric flow rate to produce a targeted, homogenously distributed shear stress on the sample

To reproduce flow characteristics like those found in the arterial vessel, high shear stress of up to 12 dyne/cm^2^ must be applied to the material sample surface. To achieve this goal, an in-stream flow guide was placed in the center of the channel to rapidly develop and accelerate the flow before passing over the sample. The position of this flow guide was simulated iteratively using Autodesk CFD 2019 to compare each shear profile produced at given flow rates. The flow guide position was simulated and optimized to provide the most homogenous distribution of shear stress across the exposed material sample. This comparative analysis concluded that a 4 mm flowguide spacing with 12 ml/min provided the best shear stress distribution on the sample surface.

To perform this comparative analysis, a simplified version of the flow channel was created using Autodesk Inventor for various flow guide positions to perform the simulations. The in-stream flow guide (Figure 2C, left side) was placed at 0-5 mm distance from the beginning of the transition to the sample surface. Preliminary simulations suggested that volumetric flow rates between 10 and 15 ml/min generate shear stress distributions that fell within the desired range. Therefore, for each model, a range of volumetric flow rates from 5 - 20 ml/min was simulated. Each combination of flow guide position and flow rate was referred to as a “scenario”. Sampling lines were used to record the shear stress at equally spaced points along each line. These lines are placed downstream of the flow guide (Figure 2A) to determine flow development (Figure 2C) and across the surface of the membrane (Figure 2D) to determine surface shear stress. The sum deviation of each of the eight sampling lines from the target value was calculated for each scenario. An example of the difference measurement for a single sampling line can be seen in Figure 2E.

### CFD simulations using a high mesh density near the region of interest show controlled flow acceleration to a homogenous shear stress distribution on the sample

Using the methodology described previously, a 4 mm flow guide spacing with 12 ml/min flow was chosen as the geometry that provided the best shear stress distribution. By introducing the flow guide, the overall shear stress could be increased, and the shear stress distribution could be modified to generate a wider range of laminar flow with high shear stress. Figure 2F illustrates the development of the flow profile between the flow guide and the center of the sample surface comparing the shear stresses along eight sampling lines with the direction of flow indicated. The shear profile clearly developed from the two accelerated streams into a smooth distribution near the chosen shear stress and the desired flow pattern was established before the flow reached the sample surface. The shear profile for the sample surface can also be seen in Figure 2F.

The histogram of simulated shear stress values for the entire sample surface is given in Figure 3C, shown in orange. 80.5% of the simulated values fell within the physiological range for arterial endothelial cells in large vessels of 8-12 dyne/cm^2^. The shear stress across the plane was quite homogenous, with 75.5% of simulated shear values within +/-1 dyne/cm^2^ of the target shear stress of 10 dyne/cm^2^.

**Figure 3.**
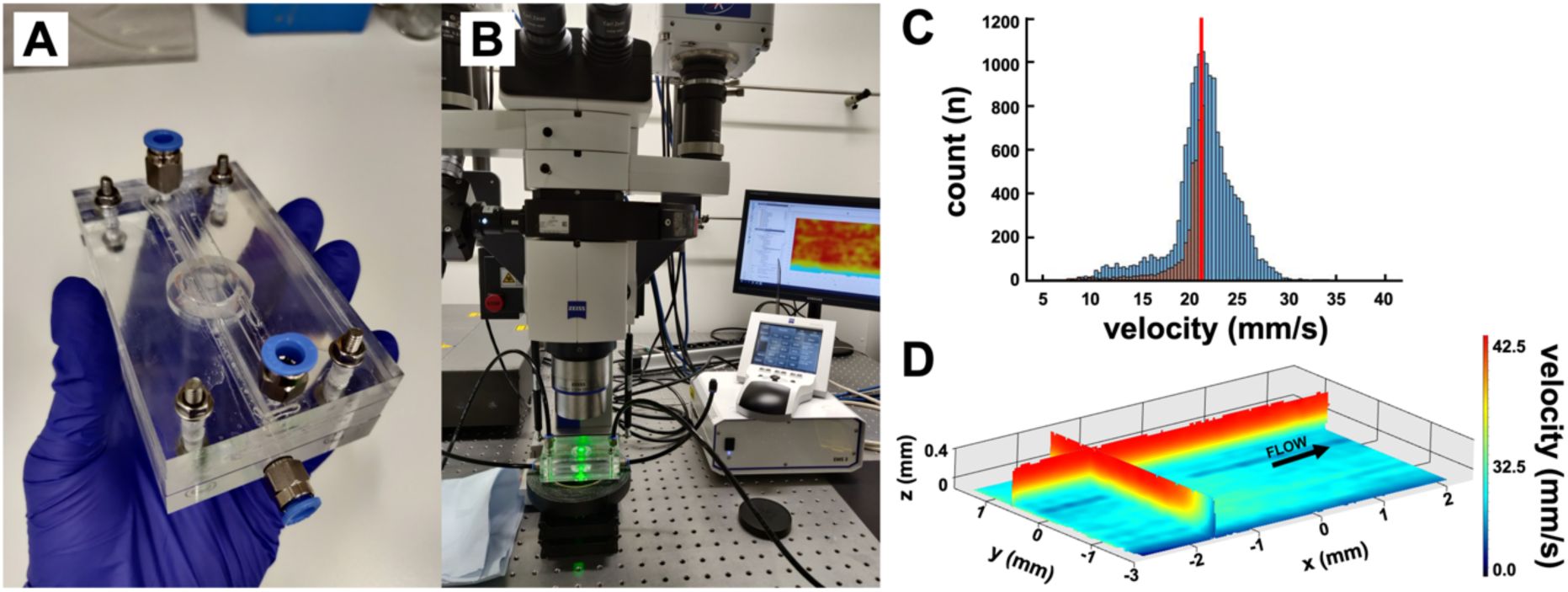
CFD simulation results validated with experimental flow analysis using microPTV. (A) Modified acrylic flow chamber designed to be compatible with microPTV equipment, assembled with tubing connectors and PCU membrane fixed inside. (B) Experimental setup of the microPTV system including the stereo microscope and the flow chamber. (C) Histogram of simulated velocities taken from CFD results (orange) superimposed over measured velocities using microPTV. Mean value of measured velocities indicated by red line. (D) Visualized microPTV results near the membrane surface using 3.2 um particles at 10 ml/min flow rate. Direction of flow indicated. The histogram of shear stresses (C) show agreement between simulated (orange) and measured (blue) values. Velocity analysis (D) indicates laminar flow across the sample surface.

### MicroPTV measured velocities compared to CFD simulated velocities

Figure 3 shows a flow chip specifically modified to fit the microPTV system, yet identical to the flow chip shown in Figure 2 in every other regard. The microPTV system in Figure 3B employed 3.2 µm polystyrene tracer particles with 10 ml/min volumetric flow rate yielding the best results for comparison with the simulated values. Figure 3C shows the measured velocities nearest to the membrane surface with the simulated velocities superimposed. Figure 3D shows the velocity in the plane closest to the membrane surface was very homogenous, applying even shear stress across the sample surface confirming the modelled flow and shear stress shown in Figure 2G.

### Calcification propensity of popular cardiac valve replacement material, bovine pericardium, as compared to polycarbonate urethane

Next, we compared calcifications generated in the large CVE-FT2 and the flow chip. Figure 4A-B show that bovine pericardium was typically heavily calcified two weeks in the CVE-FT2, while polycarbonate urethane was typically not calcified (Figure 4 E-F). Materials were also calcified in the flow chip for two weeks. As before, calcified lesions were readily detected in the pericardium (Figure 4C-D), while polycarbonate urethane was not calcified (Figure 4G-H). Strong red background staining in B and D was due to pericard autofluorescence.

**Figure 4.**
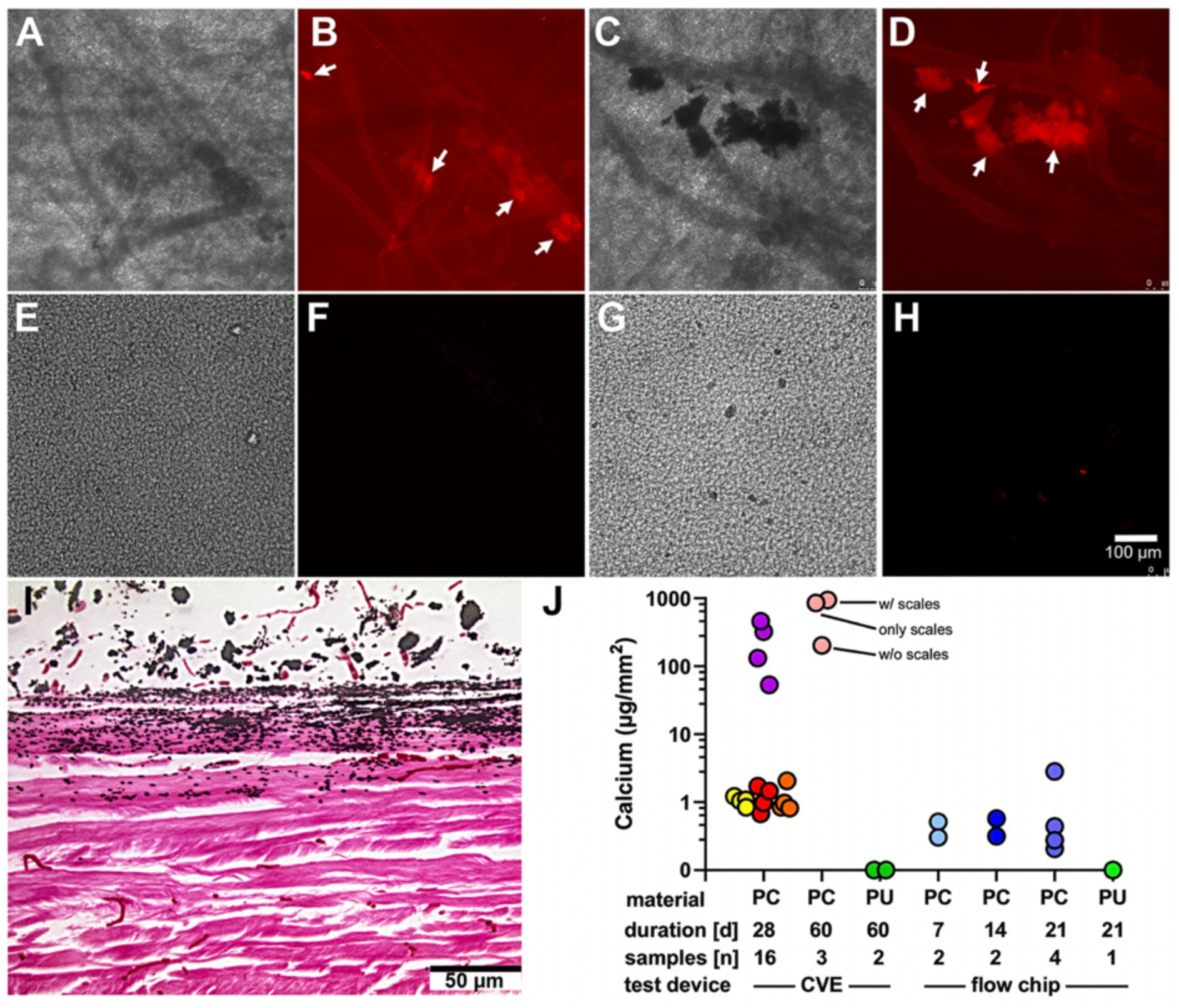
Comparison of calcifications after treatment within the CVE-FT2 or the flow chip. All samples were cultured in simulated body fluid with elevated calcium and phosphate. Samples were cultured for 14-60 days in the CVE-FT2 and 14-21 days in the flow chip. Pericard samples calcified for 14 days in both the CVE-FT2 (A,B,E,F) and the flow chip (C,D,G,H) are presented for comparison. Brightfield (A,C,E,G) and Alexa546-fetuin-A fluorescence (B,D,F,H) are shown. In both testing systems, pericard patches (A-D,I), but not polycarbonate urethane patches (E-H) stained positive with Alexa546-fetuin-A depicting calcified lesions (dark lesions in brightfield micrographs). (I) Brightfield picture of von Kossa-stained bovine pericard section after 14 days in the flow chip depicting calcified lesions on the side exposed to flowing calcification fluid (top of image). (J) Quantification of calcification using a colorimetric assay confirmed heavy calcification of pericard (PC) patches ranging from 1-1000 µg/mm^2^ and no calcification of polycarbonate urethane (PU) patches at 0 µg/mm^2^. Large variability indicates that once calcification starts, it can progress rapidly into surface scales. Each color represents one individual patch. Data points represent replicates created by splitting tested samples. CVE-FT2 patches were split in quarters and measured as technical replicates. Flow chip patches were split into two halves and measured as technical replicates. One heavily calcified PC patch incubated in the CVE-FT2 for 60 days (pink) was halved and measured with surface scaling intact (w/ scales) and separated into patch material only (w/o scales) and the scales only, creating three measured samples indicated in the figure. Total sample count per tested condition is indicated in the figure legend.

Next, we studied the calcification of pericardium in cell-compatible calcification medium in view of subsequent calcification studies on biohybrid constructs. The calcification medium (CM) was based on DMEM cell culture medium. Figure 4I shows von Kossa-stained pericardium calcifications after one week flow chip incubation in CM. Calcified lesions were present inside tissue layers of the pericardium.

The calcification can clearly be seen to have progressed from the surface, which was exposed to CM in the high-flow channel of the two-compartment chip (Figure 4I, top). The opposing surface (Figure 4I, bottom) had been exposed to DMEM basal medium (BM) in the low-flow channel of the two-compartment chip and showed no signs of calcification. The sample was calcified for 14 days only, showing that rapid calcification was possible even in physiological, cell-compatible conditions. This allowed the use of our flow chip as a test system for biohybrid constructs including cells.

### Calcium content of calcified materials

The calcium content of the tested materials was quantified using a colorimetric assay (Figure 4J). Bovine pericardium (PC) patches calcified in both systems, with the CVE-FT2 sometimes developing heavy surface scaling that led to higher calcium content variability. The flow chip produced calcification within the tissue itself without extensive surface scaling. In contrast, polycarbonate urethane (PCU) patches showed no calcification in either system. Each color in Figure 4J represents an individual patch. For the CVE-FT2, patches were split into quarters and measured as technical replicates. Flow chip patches were measured whole. One PC patch from the CVE-FT2 incubated for 60 days (pink) was cut into two halves. One half was analyzed with surface scales intact (’w scales’), and one half was cleaved into the scales (’scales only’) and the underlying patch material (’w/o scales’) demonstrating that the majority of the mineral was contained in the surface scales.

### Textile-reinforced biohybrid construct

Having validated the efficacy of cell-compatible calcification medium in the flow chip, we finally studied calcification of a cellularized biohybrid construct comprising a blended woven textile and fibrin-gel scaffold populated with vascular smooth muscle cells. After 7 days of exposure to flow in calcification medium, biohybrid constructs maintained hydrogel integrity, cell viability, yet showed clear signs of early calcification. Figure 5A shows a cellularized biohybrid construct mounted inside the assembled flow chip. Imaging was performed with the samples both inside (Figure 5B-E) and removed from the chip (Figure 5F-H). Calcifications are distinctly discernible between the cells at higher magnification (arrows in Figure 5H). Utilizing Celltracker Green staining, cell viability was demonstrated to be preserved throughout the duration of the test (Figure 5D-H). After 7 days of culture in CM, cells persisted in populating the entire bulk gel (Figure 5G-H). Gel integrity was maintained even in the regions subjected to high flow.

**Figure 5.**
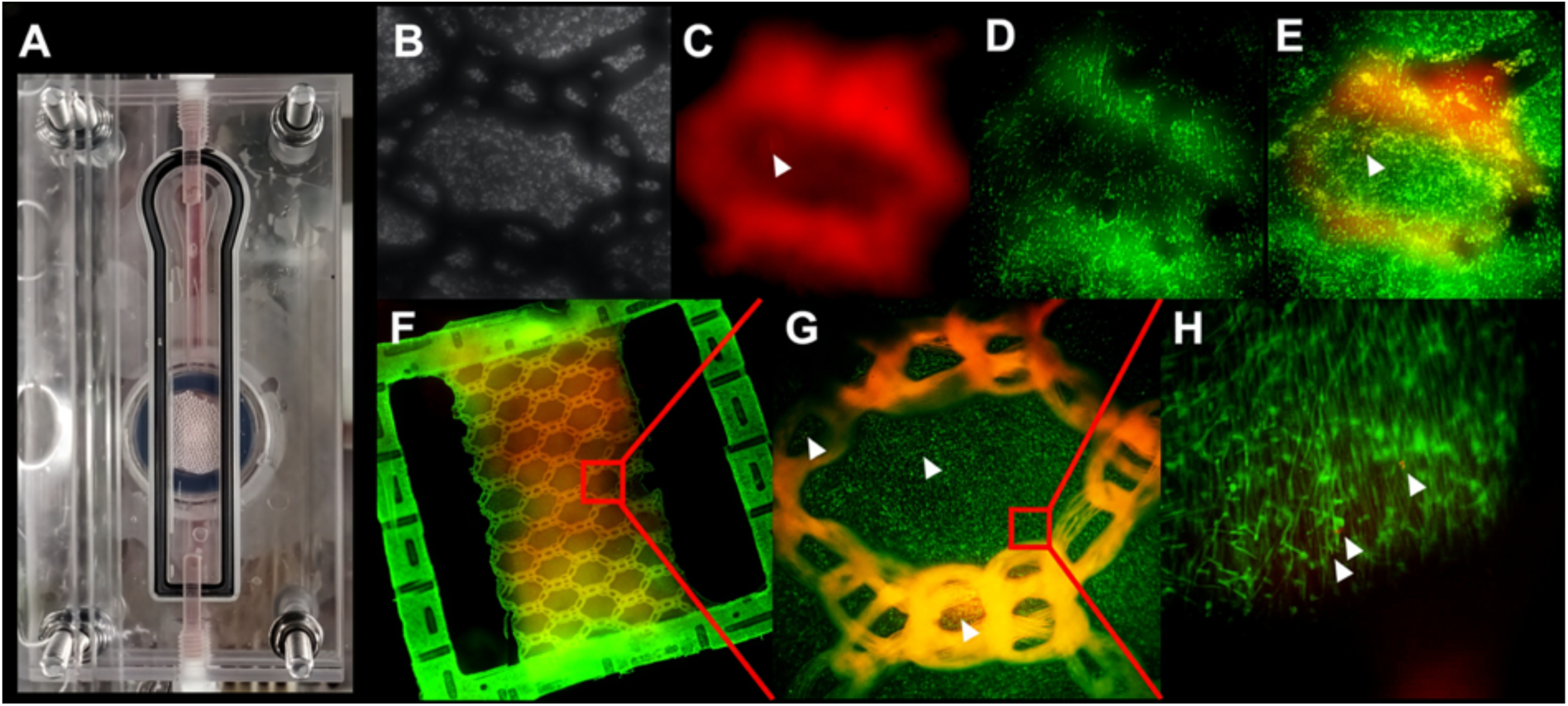
Three-piece flow chip compatible with live microscopy of calcification of biohybrid samples using fluorescent fetuin-A. (A) Assembled flow chip containing a textile-reinforced biohybrid construct with embedded smooth muscle cells. Samples were cultured under flow conditions in calcification medium for 7 days. Figures B-E show the biohybrid construct imaged while inside the flow chip. (B) Brightfield image shows the textile fiber reinforcement. (C) Red fluorescence shows textile autofluorescence as well as fluorescent fetuin-A staining for calcification marked by white arrows in C, E, G and H. (D) Green fluorescence showing viable smooth muscle cells, labelled with Celltracker Green. (E) Composite image with all three channels overlayed. Figures F-H show a framed biohybrid construct removed from the chip, for higher resolution microscopy. (F) Whole textile-reinforced biohybrid construct consisting of a structural frame and textile reinforcement (PET), a fibrin-based hydrogel, and embedded smooth muscle cells. The dense PET outer frame is autofluorescent in both green and red channels, but red fetuin-A signal can be clearly seen in the bulk gel. (G) A single textile loop section of the biohybrid construct, showing smooth muscle cells populating the bulk gel. (H) Viable smooth muscle cells (green) with calcifications (red) indicated by white arrows.

## Discussion

The need to develop a chip-based cell calcification system was driven by a lack of suitable commercially available products. While many commercial chip platforms exist, none are intended to produce the desired shear stress while facilitating live imaging of both sides of the contained sample. Some products, however, do support these conditions at lower flow rates. These devices expose contained membranes to low crossflow on both sides and can be used as a kidney model.^[32]^ One such device was tested for our purposes, but while the device performed as expected at lower flow rates, and thus lower shear stress, leaking and loss of function were observed when the flow rate was increased to generate shear stresses like those found in arterial vessels. Calcification trials were first performed with non-cellularized materials, bovine pericardium and polycarbonate urethane. The comparative study between the newly developed flow chip and the already established CVE-FT2 yielded similar calcium content for both devices in the same order of magnitude, yet heavy scaling was not developed in the flow chip (Figure 4). Fetuin-A based imaging enabled visualization of small, calcified lesions in the pericardium in both testing systems. (Figure 4B, F) However, non-destructive longitudinal imaging was only possible with the flow chip device. (Figure 1A-C) Pericardium as a bioprosthetic material is prone to extensive calcification. This calcification is commonly attributed to cell debris that are not removed during the glutaraldehyde fixation process, and to residual glutaraldehyde, which introduces nucleation sites for calcification.^[33]^ The mineral deposition further extends along the collagen and elastin fibers within the pericardium. In our results, small calcification deposits could be observed along the collagen fibers, confirming the expected mineralization pattern. (Figure 4A, E) The calcification fluid used for this comparison had already been proven to calcify bovine pericardium in the large-scale CVE-FT2 within two weeks. In the flow chip, we were able to generate comparable calcification of the material proper in a short-term test of two weeks. Nevertheless, the fluid’s application was limited to cell-free materials due to the added biocide. Using biocide was necessary in the CVE-FT2, since it was an open system. Polycarbonate urethane did not calcify after treatment with near supersaturated mineral solutions, serving as negative control in our study. Both test systems did not induce calcification in polycarbonate urethane. (Figure 4C-D, G-H)

By generating comparable results using the newly developed flow chip and the established CVE-FT2, we demonstrated the comparability of the two platforms. Furthermore, to meet the requirement for a cell-compatible test fluid suitable for biohybrid implants, we developed a calcification medium based on DMEM, a common cell culture medium. This medium also showed positive calcification of pericardium in the flow chip and thus confirmed the usability of the flow chip for calcification testing of biohybrid implants. Analysis of acid-extracted calcium revealed comparable levels of calcification in the pericardial tissue regardless of testing device. However, the pericard samples calcified on the CVE-FT2 exhibited significant surface scaling, which was not observed in the flow chip device.

Our findings on the detrimental effects of medium exchange in static culture can be interpreted considering recent biomineralization research. Schaart et al. (2025) identified calciprotein particles (CPPs)—complexes of fetuin-A, calcium, and phosphate—as stable mineral carriers in serum that prevent spontaneous precipitation. Their work suggests that early-stage CPPs can directly mineralize collagen, while matured CPPs require cellular processing for controlled deposition^[23]^. In our static cultures, each medium exchange likely introduced a bolus of fresh, early-stage CPPs. This sudden shift in mineral saturation, as reflected in our data (Figure 1A-B), appears to drive cell-independent, extracellular matrix calcification and coincides with increased cell death. Thus, the conventional static culture protocol with periodic feeding creates a non-physiological cycle of mineral supersaturation that artifacts the calcification process.

In contrast, our continuous flow system circumvents this problem by maintaining a stable, homeostatic mineral environment. The constant perfusion avoids the peaks and troughs in CPP concentration, allowing for the maintenance of stable mineral phases. This prevents the spontaneous mineralization observed after each static medium exchange and appears to enable cells to regulate mineral deposition through native endocytic and recycling pathways. The result is a more physiologically relevant model where calcification progresses steadily over time (Figure 1C), not in abrupt jumps (Figure 1B). Importantly, calcification is not prevented but is rather re-localized to areas of high shear stress within a largely viable cell population (Figure 1D-F). This demonstrates that the flow chip enables a cell-mediated calcification process, distinct from the passive precipitation dominating in static culture.

To study the calcification of biohybrid samples, a textile-reinforced fibrin-based construct populated with vascular smooth muscle cells was also calcified in the flow chip with physiological shear stress (Figure 5). After 7 days, the biohybrid constructs showed calcification and high cell viability (Figure 5 D-E) like the cells shown in Figure 1E, confirming that the absence of cell death-associated calcification caused by medium exchange did not prevent calcification altogether. Continuous live imaging of calcifications was achieved by adding fluorescent fetuin-A to the circulating medium. This serum-derived protein is a major component of fetal bovine serum anyway and therefore proven non-toxic. In addition, it has been shown to preferentially bind nascent calcium phosphate mineral thus staining calcifications as they form.^[25]^ The calcified lesions, as well as viable cells, were clearly imaged while the sample remained secured inside the flow chip (Figure 1 A-E). Upon removal from the device for higher resolution microscopy, potential calcification nucleation sites could be observed in the early stages of development in the vicinity of viable cells (Figure 1 F-H).

In terms of experimental conditions, the flow chip was advantageous over larger testing devices concerning the use of resources, especially calcification fluid and experimental material patches, as well as energy demands, hazardous substances involved, and space requirement. In addition, the flow chip minimized the required amount of imaging reagents like fluorescent fetuin-A, and allowed non-destructive, longitudinal monitoring of calcification in a standard CO_2_ incubator, present in most biological labs. This may benefit the study of a wider range of calcification pathologies associated with tissue or metabolic abnormalities, disease, or implantation of certain biomaterials.^[34]^

### Limitations

While this device represents the first steps in developing a calcification testing platform for cellularized biohybrid implant materials, there remain important factors to be considered and implemented into future work. One key factor is the inclusion of endothelial cells to the materials during calcification. Both endothelial cells and vascular smooth muscle cells must be present to study the response of the endothelium to calcification and to investigate the crosstalk between the two cell types. Culturing both cells in calcifying conditions requires the development of cell-specific and cell-compatible calcification medium. This is part of our ongoing work.

Additionally, while maintaining a homogenous and stable application of shear stress to the material surface allows for fine control over experimental conditions, it does not accurately mimic the flow conditions in the vessel lumen which is driven by the heart. In the CVE-FT2, the patches are mechanically stressed through the repetitive motion of the drive unit mimicking a beating heart. While comparable calcification results were obtained in the chip, the lack of mechanical stimulation and cardiac waveforms is noted and will be the topic of future work on this device as it was certainly relevant to the mechanisms of vascular calcification.

## Conclusions

In summary, we developed a miniaturized, two-channel flow chip capable of simultaneously exposing a cellularized material sample to two unique flow and medium conditions to investigate the mechanisms of calcification that demonstrated comparable results to those produced by and established large, yet less versatile CVE-FT2. The flow characteristics in the high-flow channel of the chip were simulated using CFD and validated using microPTV, showing that the flow was sufficiently developed and accelerated to generate physiological shear stress. These results demonstrate the comparability of this miniaturized flow chip to larger testing devices for the purpose of detecting, monitoring, and imaging calcification in real-time. The miniaturized flow chip platform offers the benefit of requiring significantly smaller material patches and volumes of fluid, as well as the novel opportunity to monitor the progression of calcification from two different environments. We believe this platform will allow for unique insights into the origin and progression of calcification-related cardiovascular disease as well as the opportunity to examine interventions in real-time.

## Author Contributions

A. Morgan: Conceptualization, Methodology, Software, Validation, Formal Analysis, Investigation, Resources, Data Curation, Writing – Original Draft, Writing – Review & Editing, Visualization, Project Administration. R. Dzhanaev: Investigation, Validation. A. Gorgels: Methodology, Investigation, Writing – Review & Editing, Visualization. L. Stüwe: Methodology, Investigation, Formal Analysis, Writing – Review & Editing. F. Stockmeier: Methodology, Investigation, Formal Analysis, Writing – Review & Editing. C. Böhm: Methodology, Investigation, Writing – Review & Editing. J. Clauser: Supervision, Writing – Review & Editing. S. Jockenhoevel: Conceptualization, Resources, Supervision, Project Administration, Funding Acquisition, Writing – Review & Editing. U. Steinseifer: Conceptualization, Resources, Supervision, Funding Acquisition, Writing – Review & Editing. W. Jahnen-Dechent: Conceptualization, Resources, Supervision, Project Administration, Funding Acquisition, Writing – Review & Editing.

All authors have reviewed and approved the final manuscript.

## Conflicts of interest

There are no conflicts to declare.

## Acknowledgements

This work received funding from the German Research Foundation (DFG, Deutsche Forschungsgemeinschaft) – TRR 219 – Project-ID 322900939 (W. Jahnen-Dechent) and 403041552 (W. Jahnen-Dechent and U. Steinseifer).

## Notes

### Competing Interest Statement

The authors have declared no competing interest.

